# MALDI-FISH for co-localization of brominated metabolites and *Pseudovibrio* spp. in *Aplysina* tissue

**DOI:** 10.64898/2025.12.19.695590

**Authors:** Allyson McAtamney, Cayla Lagousis, Bradley S. Colquitt, Lucia Pita, Alessandra S. Eustaquio, Laura M. Sanchez

## Abstract

Bromotyrosine-containing natural products have long been isolated from marine sponges of the order *Verongiida*. Recent studies have questioned the source of these natural products whether production occurs through the sponge, its diverse microbiota, or perhaps a combination of both. While studying the uptake of engineered bacterial strains by sponges, we have observed production of fistularin-3 in sponge. To explore these results further, we employed multimodal imaging techniques to co-localize microbial metabolites with fluorescent in situ hybridization (FISH) probes. Here, we built on previously developed MALDI-FISH methods to add to the conversation on brominated metabolite production in marine sponges. Using MALDI-MSI, we measured fistularin-3, a poly-brominated natural product previously isolated from *Aplysina aerophoba*, present in our *A. aerophoba* samples that were inoculated with *Pseudovibrio brasiliensis*. A recent study reported that *P. brasiliensis* produced fistularin-3 in pure culture, but this data has not yet been reproduced. We employed MALDI-MSI and FISH on the same sponge cryosections to colocalize the spatial distribution of fistularin-3 with *P. brasiliensis* in sponge tissue. Despite the caveats of this study, our data suggests that perhaps both sponge and microbe may be required for production of fistularin-3.

## Introduction

We initially sought to identify pseudovibriamides from different strains of *Pseudovibrio brasiliensis* in both bacterial colonies and *Aplysina aerophoba* sponge tissue.^9^ These secreted compounds have been shown to promote motility and reduce biofilm formation, and we hypothesized that these functional roles might provide a colonization advantage in sponge.^9^ We generated deletions of the pseudovibriamide biosynthetic gene cluster with the goal of understanding their role in host colonization. However, we discovered the presence of fistularin-3 and related brominated derivatives in *A. aerophoba* tissue using MALDI-MSI. Upon this finding, we sought to build on the previously developed methods to add to the conversation on fistularin-3 production in marine sponges.

Considering the difficulties in replicating the experiments from Nicacio *et al.*, we employed MALDI-FISH multimodal imaging to better understand the spatial production of fistularin-3, a bromotyrosine-derived alkaloid previously isolated from both *A. aerophoba* and *Pseudovibrio brasiliensis*.^8,10^ In revisiting the source of brominated natural products, another group explored a variety of bromotyrosine containing sponges, their microbiomes, and their metabolites and concluded that sponges harbor diverse microbiomes with diverse metabolomes, where the polar metabolome provided insight into potential biosynthetic precursors.^11^ Ultimately, the study concluded that the source of these brominated metabolites is unclear, but there is a strong correlation between the sponge metabolome and architecture of its microbiome.^11^ We also found that there is a strong correlation between the distribution of fistularin-3 in the sponge and *Pseudovibrio* presence, and note that presence of the compound is mainly found along the oscules and outer layer of the sponge tissues. We hypothesize that production of this metabolite may require both the sponge and this microbe.

## Methods

### Bacterial strains and cultivation

*P. brasiliensis* Ab134 wild type and Δ*pppA* mutant strains were cultivated at 30 ℃ on BD Difco™ Marine Agar 2216 (MA) or in BD Difco™ Marine Broth 2216 (MB) for 18-24 h unless otherwise noted. *P. brasiliensis* strains expressing fluorescent proteins were obtained as described previously.^12^ GFP was expressed from plasmid pAM4891 and mCherry from pSEVA237R_Pem7, respectively. Kanamycin (200 µg/mL) was used for plasmid selection when preparing seed cultures. All strains were cryo-preserved in 20% glycerol [v/v] at −80 ℃.

### Sponge collection

*Aplysina aerophoba* sponge specimens were collected by SCUBA in June 2022 off the coast of Girona, Spain, and kept in an aquarium with circulating, unfiltered seawater at the facilities in ICM-CSIC, Barcelona, Spain. Using a scalpel, sponge individuals were divided into specimens containing only one osculum while maintaining them submerged in seawater. Each specimen was affixed to a plastic support using a metal pin and kept in the aquarium with circulating, unfiltered seawater until use (Figure S1A-C). Sponge specimens were used within three weeks of collection.

Preparation of *P. brasiliensis* cells for sponge-bacteria studies.

*P. brasiliensis* Ab134 wild type and Δ*pppA* mutant cultures containing either pSEVA237R_Pem7 (mCherry)^13^ or pAM4891 (GFP)^14^ were initiated by inoculating 0.2 mL cryopreserved cultures onto 20 mL of marine broth with kanamycin (200 µg/ml final concentration). The cultures contained in 250 mL Erlenmeyer flasks were incubated at 27 ℃, 200 rpm, for 23 h. Cells were harvested by centrifugation (10 min at 25 ℃ and 3,000 rpm = 1,730 g). After discarding the supernatant, cells pellets were washed twice, each time with 20 mL seawater (same centrifugation conditions as above). After the last wash, the cell pellets were resuspended in 20 mL fresh seawater. An aliquot of each cell suspension was diluted 1:1,000 with seawater and cell count was performed using flow cytometry (CytoFLEX S, Beckman Coulter) set at medium flow (30 µL/min), with data acquisition for 2 min per sample. A seawater sample was used as “blank” to correct the total cell count. Wild type and mutant strains expressing either GFP or mCherry, respectively (or vice versa) were mixed at 1:1 ratio (5 x 10^8^ cells of each per mL) to yield a total of 1 x 10^9^ cells per mL. The mixed cell suspensions were kept on ice until use.

### Sponge-bacteria studies

Four experimental aquaria were prepared, each containing 4 L of filtered (50 µm) seawater and an aquarium pump. *A. aerophoba* sponge specimens containing one osculum were transferred to two of the experimental aquaria, whereas the other two aquaria remained as no sponge controls. The sponge specimens were transferred using a plastic bag, taking care that they were always submerged in seawater. Aliquots (1.62 mL) were taken from each aquarium before inoculation (time −1). 4 mL of each bacterial cell mixture (wild type expressing GFP and Δ*pppA* mutant expressing mCherry or wild type expressing mCherry and Δ*pppA* mutant expressing GFP) was then added to two aquaria (to make up 1 x 10^6^ cells per ml final concentration), one of which contained a sponge specimen. Water samples (1.62 mL) were taken from each aquarium at 0, 0.5, 1, 2, 3, and 4 hours, transferred to 2 mL Eppendorf tubes and mixed with 0.18 mL fixative (10% paraformaldehyde, 0.5% glutaraldehyde). After 10 min incubation at room temperature, the fixed samples were stored at −20 ℃. Flow cytometry was performed on all samples as described above to determine cell count originating from cells expressing mCherry and GFP. The mCherry and GFP gates were set based on pure cultures. The relative cell count for each aquarium was calculated after correcting for the corresponding −1 (before inoculation) sample to yield the results shown in Figure S2.

MALDI-MSI sample preparation of *P. brasiliensis*.

Overnight liquid cultures in marine broth (BD Difco, Fisher) were diluted to an OD_600_ of 0.1, and 5 µL of culture was spotted onto 100 mm petri dishes with 20 mL marine agar (BD Difco, Fisher). Colonies were grown for 2 days then excised from Petri dishes using a razor blade and transferred to a ground steel MSP target plate. Using a 53 µm stainless steel sieve (Hogentogler, Inc.), colonies were covered with recrystallized matrix (1:1 dihydroxybenzoic acid (DHB): α-cyano-4-hydroxycinnamic acid (CHCA)) and dried in a 37 °C incubator and spun using a plate spinner for one hour. Following the application of a second layer of matrix, colonies were dried at 37 °C for at least 2 more hours or until dry, for a total of at least 3 hours of drying time. The excess matrix was removed using a gentle stream of air. Phosphorus red (supersaturated in MeOH, Sigma) was used as a standard and spotted onto a clean area of the plate and air dried.

MALDI-MSI sample preparation of *A. aerophoba*.

*A. aerophoba* samples were cryosectioned using a Leica CM3050S at 50 µm and thaw mounted onto a glass slide (Fisher). Bacteria (1 µL of an overnight culture) of the follow species was spotted away from the sponge samples and air dried: *P. brasiliensis* WT mCherry, *Vibrio fischeri* MJM 2386, *Vibrio cholerae* smooth, *Pseudomonas aeruginosa* PA01, *Bacillus subtilis* 3610, and *Escherichia coli* K12. α-Cyano-4-hydroxycinnamic acid (CHCA) matrix was applied using a HTX matrix sprayer. CHCA was prepared at 10 mg/mL in 9:1 ACN:H_2_O and sprayed onto the sample with a nozzle temperature of 70 ℃ and 12 passes, resulting in a matrix density of 0.007273 mg/mm^2^. 1 µL of phosphorus red (supersaturated in MeOH, Sigma) was spotted onto the glass slide and used as a calibrant.

### MALDI-MSI instrumentation

All MALDI-MSI data was acquired on a Bruker timsTOF FleX QqTOF mass spectrometer (Bruker GmbH, Billerica, MA). Imaging mass spectrometry data was acquired using timsControl v. 6.0.0 and flexImaging software v. 7.5 (Bruker Daltonics, Billerica, MA). Imaging data was acquired in positive mode from 100 to 1250 Da with a 50 μm M5 defocus laser. All step and spot sizes for sponge experiments (S) were acquired at 50 μm (50 × 50 μm laser size), and the mass resolution (R) in all experiments was 65,000 FWHM at *m/z* 1222. Bacterial imaging experiments in **Figure 1** were acquired with a step and spot size (S) of 200 μm (200 × 200 μm laser size). Other details regarding the acquisition parameters, including scan number/shot count/frequency (S), molecular identification (M), annotations (A), and time of acquisition (T) are indicated in each figure caption using the “SMART” standardized nomenclature.^15^

**Figure 1.**
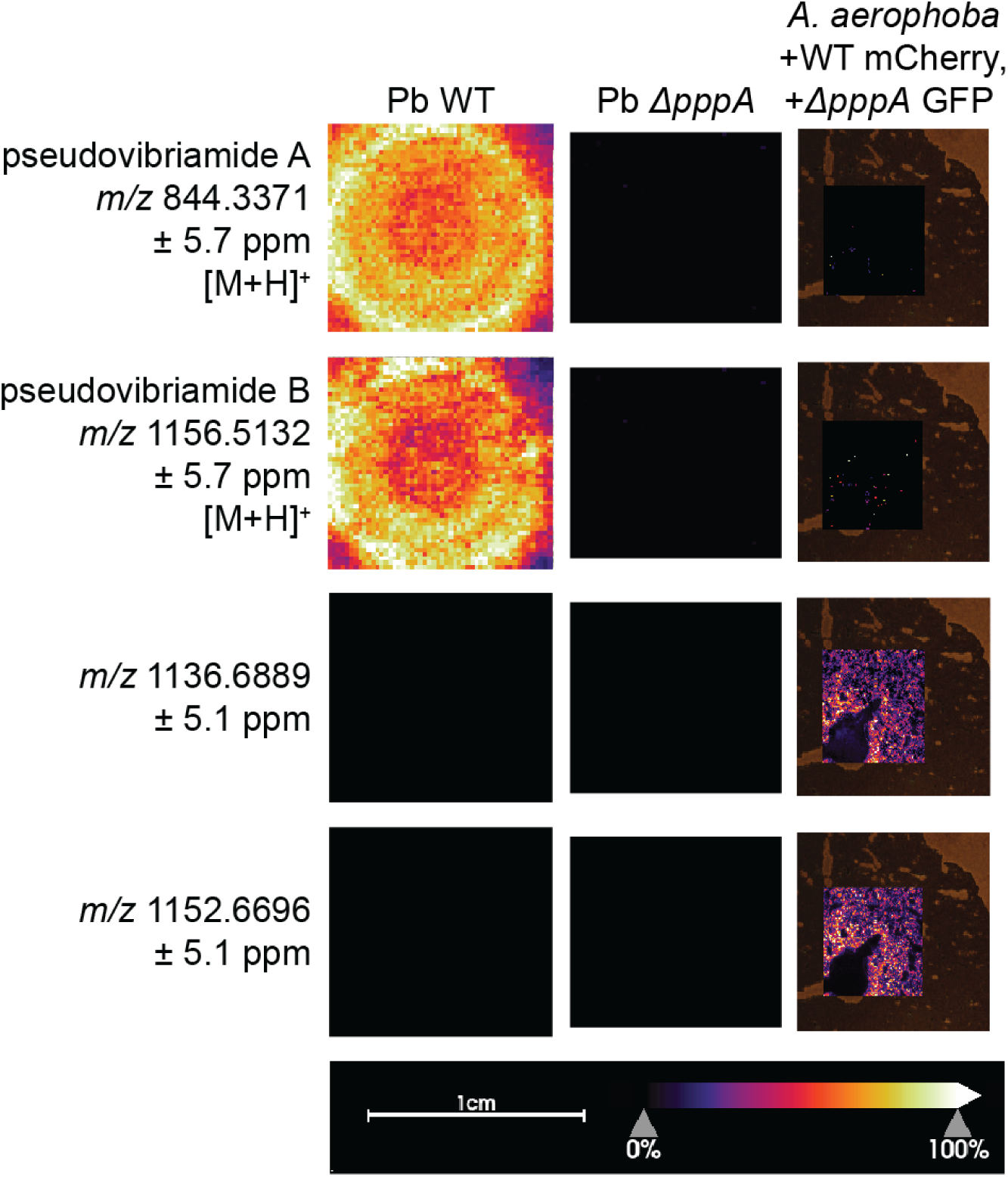
MALDI-MSI on *P. brasiliensis* colonies on agar and sponge thin tissue section. Both *P. brasiliensis* WT and Δ*pppA* mutant are shown. Only *P. brasiliensis* WT was observed to have signals for the pseudovibriamides. There was no signal observed in either colony for fistularin-3. There was no significant signal observed for the pseudovibriamides in sponge tissue. Signal was observed for fistularin-3 at both [M+Na]^+^ and [M+K]^+^ adducts. Bacterial MSI dataset contains S) 11,342 scans and were acquired at 500 shots and 1000 Hz with a 200 µm x 200 µm step and spot size. Pseudovibriamides M) were validated at the MS^1^ level. All annotations A) were targeted, and the total time of acquisition T) was 1 hrs, 49 mins. Sponge MSI dataset contains S) 24.294 scans and was acquired at 500 shots and 2000 Hz. Fistularin-3 M) was validated at the MS^2^ level. All annotations A) were untargeted, and the total time of acquisition T) was 1 hrs, 55 mins.

### MALDI-MSI data analysis

All ion images were generated using SCiLS lab software version 2025b (Bruker Daltonics, Billerica, MA). All ion images have hot spot removal applied and were normalized using a total ion count (TIC) normalization in SCiLS. Molecular identifications (M) were done to the MS^2^ level, all annotations (A) were untargeted, and GNPS2^16,17^ and CHEMnetBASE Dictionary of Natural Products 34.1 were used to make annotations. Information regarding these annotations are indicated in the figures.

### MALDI MS/MS

All MALDI-MS/MS data was acquired on a Bruker timsTOF FleX QqTOF mass spectrometer (Bruker GmbH, Billerica, MA) using Collision-Induced Dissociation (CID) via N_2_ gas. All collision energies for MS/MS spectra are provided in the figures, while isolation width was 1 Da for all spectra.

### FISH sample preparation

Following MALDI-MSI acquisition, samples containing CHCA were fixed on microscope slides using ice cold 4% paraformaldehyde (PFA)/phosphate-buffered saline (PBS) for 15 minutes. HCR probes, buffers, and hairpins were purchased from Molecular Instruments, except Probe Wash Buffer (PWB). PWB contains 50% Formamide, 25% 20× SSC, 9% 100mM Citric Acid (pH 6.0), 0.001% Tween 20, 0.5% Heparin (10mg/mL initial concentration), and 15% dH2O. The tissue was then washed three times in 5 minute increments in PBS + 0.1% Tween 20 (PBST), on a shaker. Dehydration of the tissue was performed by a series of ethanol (EtOH) washes on a shaker at room temperature (RT): 1 × 3 minute wash of 50%EtOH; 1 × 3 minute wash of 70%EtOH; 2 × 3 minute washes of 100% EtOH. The tissue was allowed to air dry for 3 minutes at RT. Next, the tissue was incubated with 37°C 150µL probe hybridization buffer before 4nM of each initiator probe (final concentration) was applied in 37°C 150µL probe hybridization buffer. The tissue was then incubated at 37°C overnight. The next day, tissue was submerged in PWB for 5 minutes at RT to allow for the release of coverslips from the tissue.

Next, the tissue was washed through a series of PWB/SSCT (5× sodium chloride sodium citrate + 0.1% Tween 20) for 15 minutes per wash at 37°C in the following order: 75%PWB/25%SSCT, 50%PWB/50%SSCT, 25%PWB/75%SSCT, 100% SSCT. The tissue was then washed once more with SSCT for 5 minutes at RT before RT 200 µL amplification buffer was applied and the tissue was incubated at RT for 30 minutes. Fluorophores were incubated at 95°C for 90 seconds and left to rest in the dark for 30 minutes at RT before being applied to the tissue in 110 µL amplification buffer. The tissue was incubated in the dark at RT overnight. Finally, on the third day, the tissue was added to 5× SSCT at RT to allow for the removal of the coverslips and then 200 µL of 5× SSCT + DAPI (1 µg/mL final concentration) was added to then incubate at RT in the dark for 30 minutes. Next, the tissue was washed for 30 minutes in 5× SSCT at RT in the dark before one more wash with 5× SSCT for 5 minutes at RT in the dark. Prolong Glass Antifade Medium (Thermofisher) was applied to each slide and coverslipped.

### FISH instrumentation

Imaging was performed on a Zeiss 880 Confocal at the UCSC Microscopy Core facility using a 10X objective, using a tiling method as needed per the size of the section. Channels 488, 546, and 647 were used to image GFP, 16S rRNA, and mCherry, respectively.

## Results and Discussion

We first sought to evaluate the ability of *P. brasiliensis* strains that do or do not produce pseudovibriamides ^9,12,18^ to be uptaken by *A. aerophoba* sponges. Sponges were challenged with a mixture of wild-type and Δ*pppA* mutant (the biosynthesis of pseudovibriamide A and B is blocked) strains expressing either GFP or mCherry (and vice versa to account for any potential fluorescent protein or vector effects). Control aquaria contained no sponge (**Figure S2A**). Over the course of 4h, the bacteria cell counts either stayed the same or increased in the no-sponge aquaria(**Figure S2B**). The increase is associated with GFP and likely due to background contamination of bacteria with natural green fluorescence (**Figure S3**). In contrast, the cell counts decreased in the sponge aquaria at an equivalent rate for wild-type and Δ*pppA* mutant strains, indicating that the sponge takes up each strain indiscriminately under the conditions tested. It is important to note that the filtering capacity varied for each sponge individual (e.g., a lower cell count was reached for experiment 2 compared to experiment 1), (**Figure S2B**), stressing the importance of performing experiments with wild-type and mutant bacterial strains in the same aquarium to allow for rigorous conclusions.

Matrix-assisted laser desorption ionization-mass spectrometry imaging (MALSI-MSI) has become a robust method to detect metabolites in a variety of biological samples, whilst providing spatial information for localization of various compounds.^19^ Combining MALDI-MSI in multimodal imaging techniques can provide spatially driven biological context of metabolite production. Bacterial autofluorescence (e.g. for cyanobacteria) and fluorescence *in-situ* hybridization (FISH) allows localizing the bacteria within the tissue and correlating their distribution with that of secondary metabolites detected by MALDI-MS. These combined multimodal imaging studies have been fruitful in demonstrating that bacteria were localized to the pores, chambers, and exterior of the marine sponge, confirming the bacteria colonize the sponge in areas directly in contact with the surrounding sea water.^20^ Additionally, MALDI-MSI showed that the secondary metabolites were localized to the same regions as the bacteria, further suggesting the sponge host serves as a mechanism to culture unculturable bacteria with complex biosynthetic machinery. MALDI-FISH specifically was critical in the detection of microbially-produced metabolites in a deep-sea mussel tissue that were otherwise not possible to cultivate in the laboratory environment, highlighting multimodal imaging a critical tool for *in situ* investigations.^21,22^

We employed MALDI-MSI to show the presence of pseudovibriamides A and B in *P. brasiliensis* WT colonies, while we observed no signal for any pseudovibriamides in the Δ*pppA* mutant (**Figure 1**). However, despite some weak signal, we were not able to confidently observe the presence of the pseudovibriamides in the sponge tissue (**Figure 1**). Despite these findings, we did observe a signal for *m/z* 1152.6, which localized around the oscule (**Figure 2A**).

**Figure 2.**
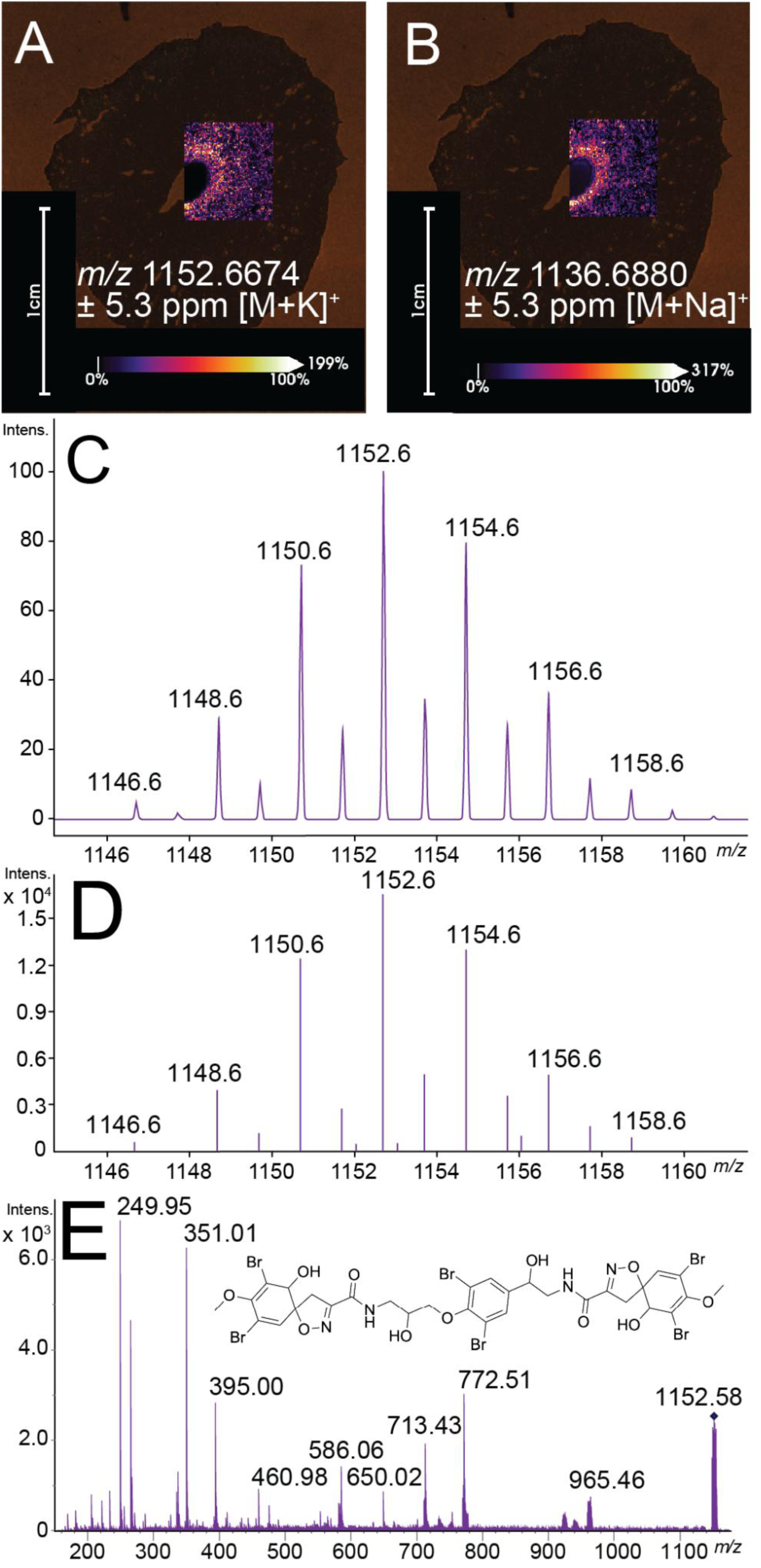
Fistularin-3 MALDI-MS spectra. **(A)** MALDI-MSI ion image of fistularin-3 [M+K]^+^ adduct and **(B)** [M+Na]^+^ adduct. MALDI-MSI parameters shown below ion images. **(C)** MALDI-MSI spectrum showing full isotope window of fistularin-3 potassium adduct. **(D)** Predicted isotopologue peaks for fistularin-3 potassium adduct using Bruker IsotopePattern. **(E)** MALDI-MS/MS spectrum of fistularin-3 obtained from the imaged sponge samples. MSI dataset contains S) 41,525 scans and were acquired at 500 shots and 2000 Hz. Fistularin-3 M) was validated at the MS^2^ level. All annotations A) were untargeted, and the total time of acquisition T) was 3 hrs, 16 mins.

The isotope distribution of *m/z* 1152.6 resembled that of a poly-brominated compound, specifically the incorporation of six bromine atoms (**Figure 2D**). Following this observation, *m/z* 1136.7 was noted to have a very similar spatial distribution around the oscule of the sponge (**Figure 2B**), and a database search in the Dictionary of Natural Products (DNP) allowed for a putative MS^1^-level annotation of fistularin-3, which was previously found to be produced by *P. brasiliensis*^8^ as well as being isolated from *Aplysina* sponges.^10^ However, we did not observe this signal directly from bacterial colonies alone (**Figure 1**). Using the chemical formula, we employed IsotopePattern software to simulate the isotopologue peaks for the potassiated adduct of fistularin-3 (**Figure 2C**), and we observed the simulated spectrum agrees with our observed spectrum (**Figure 2D**). Using MALDI-MS/MS, we obtained MS/MS spectra directly from the imaging sample and were able to confidently identify this compound as fistularin-3 at the MS^2^ level (**Figure 2E**).

Following these initial experiments, we sought to better understand the ability of either organism to produce fistularin-3, as this compound has previously been isolated from both *P. brasiliensis* bacteria, as well as *Aplysina* sponges. To do this, we employed hybridization chain reaction (HCR™) RNA fluorescent in situ hybridization (FISH) probes for imaging mRNA within a biological tissue. Targeting the 16S rRNA of *Pseudovibrio* and mCherry (**Table S1-S2**), we performed MALDI and FISH on the same thin tissue section to get accurate spatial distribution and co-localization, building off of a previously described method.^21^ Of note, the order of analysis is important to retaining signal–prior work conducted MSI then FISH, which we also followed to maximize ion image signals. We employed this method to three sponge sections, each inoculated with a different set of *P. brasiliensis* mutants: sample A contained *A. aerophoba* exposed to *P. brasiliensis* WT-mCherry and Δ*pppA* GFP; sample B contained *A. aerophoba* exposed to *P. brasiliensis* WT-GFP and Δ*pppA* mCherry; while sample C contained *A. aerophoba* exposed to *P. brasiliensis* WT-GFP only. In all three samples, colocalization of the 16S rRNA FISH probe with the spatial distribution of fistularin-3 around the oscule was observed, as well as at the outside wall of the sponge (**Figure 3**). All MALDI-MSI replicates are located in **Figure S4**. Although the oscule is where the sponge filters its surroundings, the outside of the sponge is in full contact with the surrounding marine environment, likely supporting colonization by surrounding microbes. In further examination of the 16S rRNA sequencing data for the *Aplysina aerophoba* metagenome, we indeed found more than 500 hits with very good scores (E-value < 1E-100) for the *Pseudovibrio brasiliensis* 16S rRNA.^23^ Due to the presence of *P. brasiliensis* in the available metagenome of *A. aerophoba,* we are not able to disambiguate between naturally occurring versus inoculated *Pseudovibrio* spp. Thus, fundamentally we have no negative control as a limitation of our study. This colocalization of the bacteria and compound in sponge tissue (**Figure 3**) along with the metagenomic analysis provides an intriguing correlation between fistularin and this bacterium in sponge.

**Figure 3.**
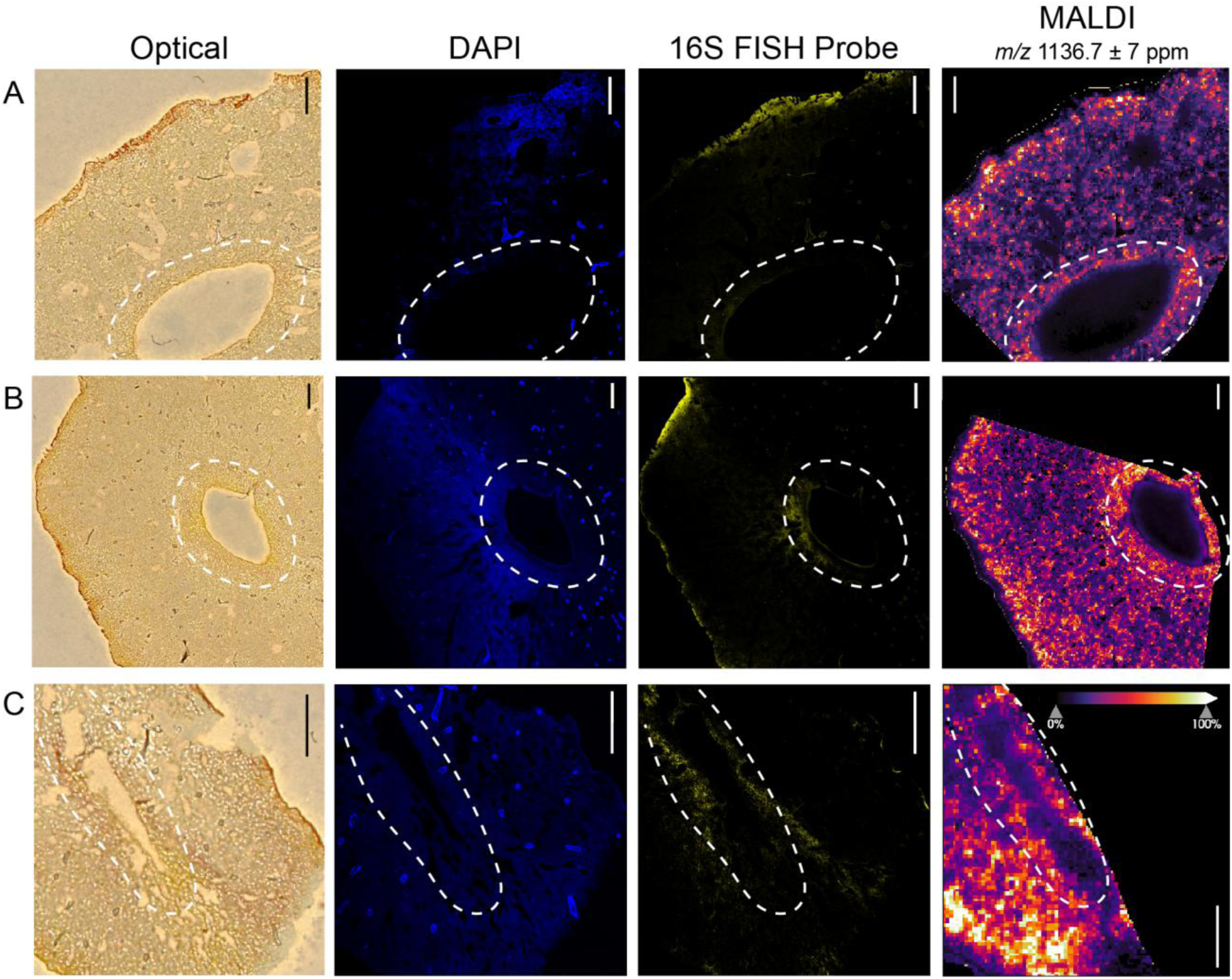
MALDI-FISH images of *A. aerophoba* cryosectioned tissue. White dashed line is placed around the oscula in each image. (A) *A. aerophoba* + *P. brasiliensis* WT mCherry, Δ*pppA* GFP. (B) *A. aerophoba* + *P. brasiliensis* WT GFP, Δ*pppA* mCherry. (C) *A. aerophoba* + *P. brasiliensis* WT GFP. MALDI-MSI parameters are as follows: S) 77,841 scans, 1000 shots, 1000 Hz. M) fistularin-3, MS^2^. A) untargeted. T) 22 hrs, 55 mins. All scale bars are 100 µm.

Furthermore, we aimed to identify additional brominated metabolites that colocalize to the same areas of the sponge section–the oscule and the outside of the sponge. MALDI-MSI data has been previously reported in this sponge, along with several brominated signals at the MS^1^ level.^7^ We observed a number of signals within a similar mass range; however, several of our observed signals have not been previously reported. Several putative brominated metabolites were identified in this study at the MS^1^ level (**Figure S5**), and we were able to identify several other brominated compounds at the MS^2^ level (**Figure 4**). For each annotation, we have included the MALDI-MSI ion image, the MS^1^ isotope pattern, and the MALDI-MS/MS spectra acquired on-sample. 3,5-dibromo-2-hydroxy-4-methoxyphenylacetamide (**Figure 4A-C**) and aplysfistularine (**Figure 4G-I**) were previously isolated from another sponge of the *Aplysina* genus.^24^ However, this study marks the first recorded detection of 3,5-dibromo-1-ethoxy-4-oxo-2,5-cyclohexadiene-1-acetamide (**Figure 4D-F**) and pseudoceratinine B (**Figure 4J-L)** in a sponge of the *Aplysina* genus, as they were previously isolated from *Suberea* spp and *Pseudoceratina* spp, respectively.^25,26^ At the MS^1^ level, we were also able to putatively identify *m/z* 258 as stryphnusin, a singly brominated phenol with acetylcholinesterase inhibition activity that has not been previously observed in the *Aplysina* genus (**Figure S5**).^27^

**Figure 4.**
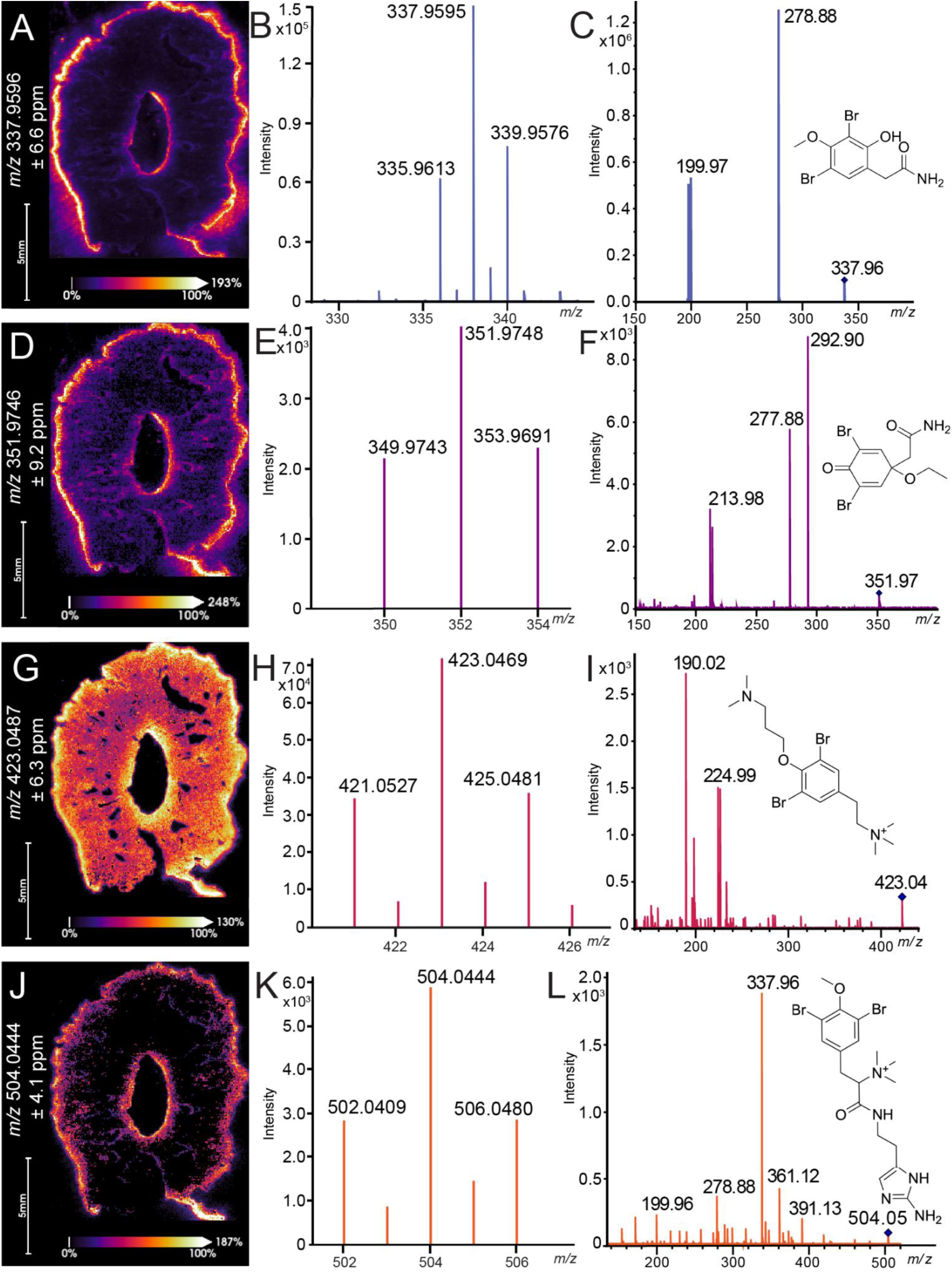
Additional brominated compounds identified in this dataset. 3,5-Dibromo-2-hydroxy-4-methoxyphenylacetamide **(A)** ion image, **(B)** MALDI-MS spectrum, **(C)** MALDI-MS/MS spectrum. 3,5-Dibromo-1-ethoxy-4-oxo-2,5-cyclohexadiene-1-acetamide **(D)** ion image, **(E)** MALDI-MS spectrum, **(F)** MALDI-MS/MS spectrum. Aplysfistularine **(G)** ion image, **(H)** MALDI-MS spectrum, **(I)** MALDI-MS/MS spectrum. Pseudoceratinine B **(J)** ion image, **(K)** MALDI-MS spectrum, **(L)** MALDI-MS/MS spectrum. Images contain S) 118,150 scans and were acquired at 500 shots and 2000 Hz. All molecular identifications M) were done at the MS^2^ level. All annotations A) were untargeted, and the total time of acquisition T) was 8 hrs, 43 mins.

In order to rule out any bacterial production of these compounds, as well as any potential FISH probe overlap, we used several overnight cultures by spotting the bacteria on the glass slide as controls. *P. brasiliensis* WT expressing mCherry was used as a positive control for confirmation of probe functionality, and the following organisms as negative controls: *Vibrio. fischeri* MJM 2386^28^, *V. cholerae* smooth^29^, *Pseudomonas aeruginosa* PA01^30^, *Bacillus subtilis* 3610^31^, and *E. coli* K12^32^. All samples that showed any signal on the confocal are shown in **Figure S6**. We did not see any signal for *P. aeruginosa* PA01, *B. subtilis* 3610, and *E. coli* K12. We used *P. brasiliensis* WT mCherry as a positive control and observed signal for the 16S FISH probe, as well as the mCherry FISH probe, suggesting the probes identified the intended target.

However, we noted that there is some signal present in *V. fischeri* for the *Pseudovibrio* 16S FISH probe, which indicates some slight level of promiscuity. However, there is no signal for the 16S FISH probe in any other organism that we tested other than *P. brasiliensis*. Additionally, there is a signal for the mCherry FISH probe in both *Vibrio* species, suggesting that there is promiscuity in that probe as well.

Given the constraints of cultivating sponges in controlled laboratory settings, these sponges were wild harvested for experiments in the lab. These sponges were environmental samples, and we cannot confirm the presence or absence of any environmental microbial strains that may exist in the natural environment, including the *Vibrio* spp. Additionally, autofluorescence may be playing a role in the mCherry probe data. We alternatively tried to image these samples through the GFP channel, which we did not have a FISH probe for.

However, we did not observe any GFP signal in our positive control, *P. brasiliensis* WT expressing GFP, suggesting the potential degradation of GFP (**Figure S7**). Although we observed mCherry signal for the *P. brasiliensis* expressing mCherry, we also saw presence of GFP signal, further suggesting autofluorescence playing a role in the GFP channel (**Figure S7**).

## Conclusion

Despite the caveats outlined above, we showed that fistularin-3 and other brominated metabolites colocalize with *P. brasiliensis* in sponge tissue, suggesting that perhaps both organisms are required for biosynthesis of these compounds. Future studies are necessary to provide more context into the production of these metabolites by each organism and the dependencies these organisms have on each other to produce potential precursors. Additionally, developing methods to culture marine sponge hosts in the laboratory environment could enhance our understanding of the interactions between marine sponges and their microbiome.^33^ Experiments that involve hosts and engineered bacterial strains could help resolve how some of these historic compounds are produced to reinvigorate investigations into their biotechnology applications.

## Acknowledgements

This work was supported by the National Science Foundation grants IOS-2220510 (LMS) and IOS-1917492 (ASE). This work was supported by the National Institutes of Health (NIH) National Institute of Neurological Disorders and Stroke NIH award R01NS138781 (BC). We gratefully acknowledge Dr. Ethan Older for helpful discussions related to NCBI queries and host-microbe considerations.

## Data Availability

All MALDI data in this study are available under CC0 1.0 Universal License as Bruker data (.d) and Bruker MALDI-MSI data (.mis) formats. MALDI-MS/MS spectra are available as open source (.mzML) format. MassIVE accession: MSV000100024.

## Supporting Information

The Supporting Information is available free of charge at https://pubs.acs.org/doi/XXX.

- ● Marine sponge culture conditions, strain table, colonization experiment details, flow cytometry data, FISH gene sequences, additional MALDI-MSI experiments, and additional confocal images highlighted in this manuscript.

