## Supplemental Information for "MALDI-FISH for co-localization of brominated metabolites and *Pseudovibrio* spp. in *Aplysina* tissue"

**Figure S1.** Tools for marine sponge-bacteria studies........................................ S2
**Figure S2.** *A. aerophoba-P. brasiliensis* colonization experiments.......................... S3
**Figure S3.** Example of flow cytometry data............................................... S4
**Figure S4.** Additional MALDI-MSI fistularin-3 replicates.................................. S6
**Figure S5.** Additional brominated MALDI-MSI signals...................................... S7
**Figure S6.** Confocal FISH images of bacterial spots....................................... S8
**Figure S7.** Additional confocal images of *A. aerophoba* and *P. brasiliensis*............... S9

**Materials and Methods:**

**Table S1.** Bacterial strains used in this study.............................................. S10
**Table S2.** Gene sequences used for HCR-RNA-FISH probe design........................... S11
**References** ................................................................................... S13

**Tools for *P. brasiliensis* bacteria-sponge studies.** *A. aerophoba* marine sponge specimens from the Mediterranean Sea, a sponge species from which *Pseudovibrio* containing the *ppp* gene cluster have been previously isolated,[1] were collected by SCUBA. The field-collected sponges were kept in aquaria with unfiltered, circulating seawater where they can be maintained for several months (Figure S1A-3C). *Pseudovibrio* wild type and Δ*pppA* mutant expressing either green (GFP) or red (mCherry) fluorescent protein encoded in plasmids pAM4891 and pSEVA237R_Pem7, respectively, were previously obtained (Figure 1D).[2]

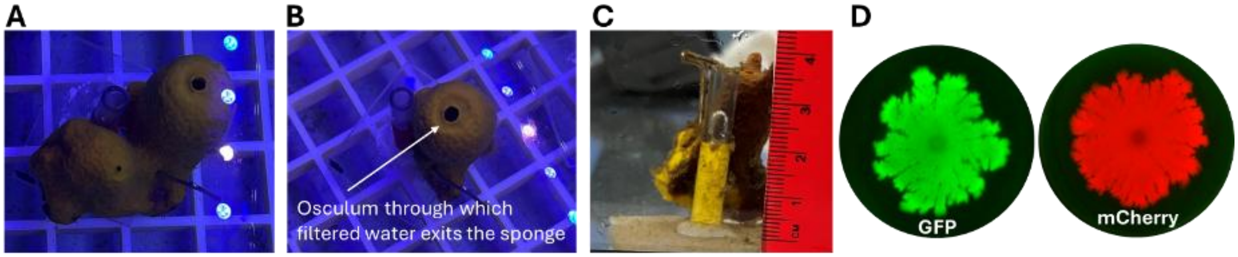


**Figure S1.** Tools for marine sponge-bacteria studies. (**A**) *A. aerophoba* marine sponge specimens were collected by SCUBA and kept in aquaria with unfiltered, circulating seawater until use. Sponges can be kept as such for several months. (**B**) Single individuals containing one osculum were sectioned using a scalpel and attached to a plastic support. These single-osculum individuals were used in sponge-bacteria experiments. (**C**) Individuals used were about 3 cm long but can vary in shape and filtering capacity. (**D**) Fluorescence imaging of strains of *P. brasiliensis* Ab134 that express green or red fluorescent proteins (GFP and mCherry) from pAM4891 and pSEVA237R_Pem7, respectively (select swarming pictures of the *pppA* mutant at 72 hours are shown). The two-color labels allow competition experiments to be performed in the same aquarium which is important to account for sponge individual variability.

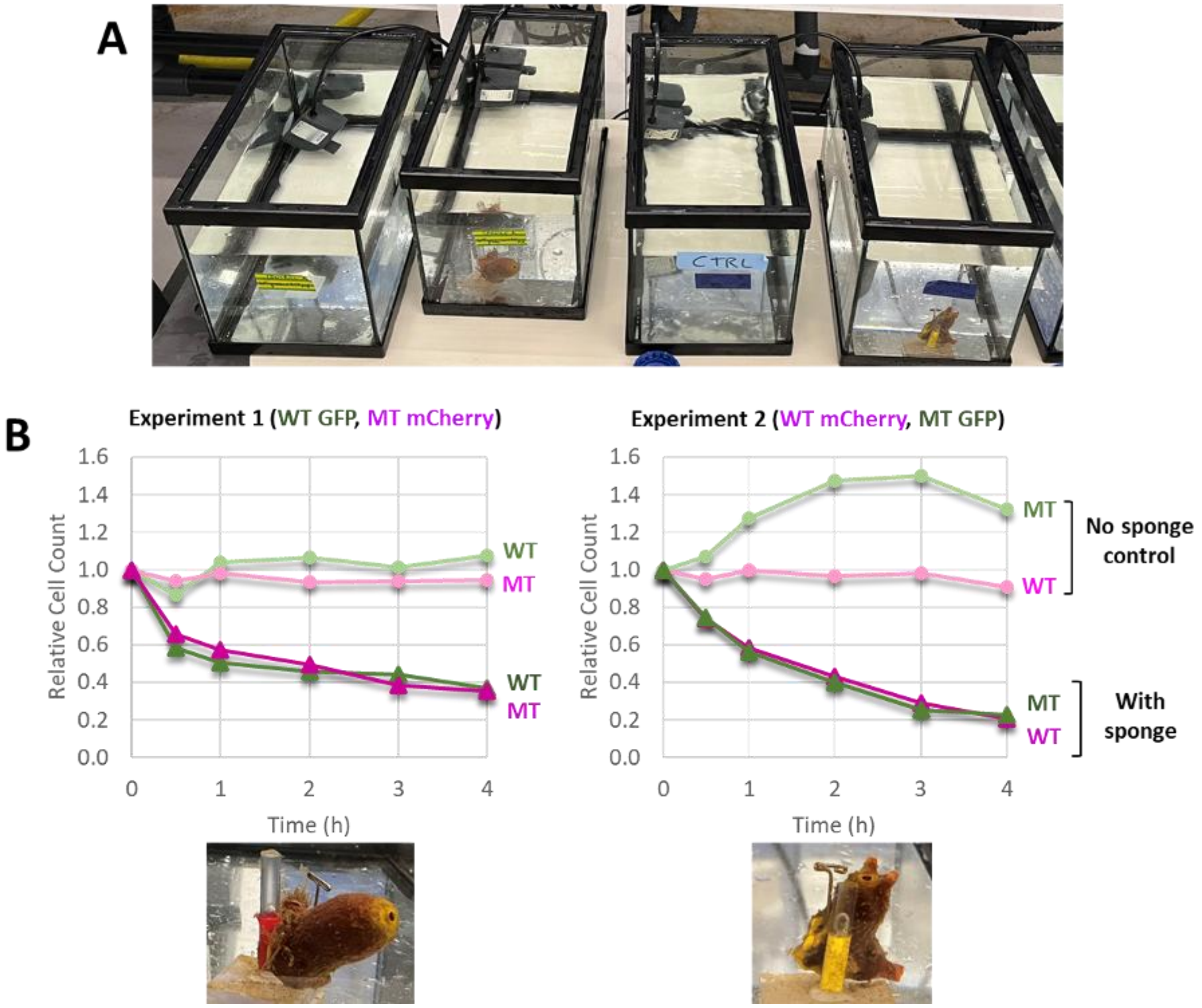


**Figure S2**. *A. aerophoba* marine sponge-*P. brasiliensis* bacteria experiments. (**A**) Four aquaria were set up with 50-µm filtered seawater, two of which contained sponge individuals, and two of which served as no-sponge control. A mixture of wild-type (WT) and Δ*pppA* mutant (MT) expressing either green (GFP) or red (mCherry) fluorescent protein were added to each aquarium at a final concentration of $10^6$ cells/mL as measured by flow cytometry. In Experiment 1, aquaria contained WT with GFP and MT with mCherry, and the labels were inverted for Experiment 2. (**B**) Relative cell count over time for each experiment as measured by flow cytometry. The sponge individuals used in each experiment are shown at the bottom. The increase in green fluorescence signal in aquaria without sponges (particularly marked for experiment 2) is likely due to seawater contamination as the sponge specimen was transferred from the long-term aquarium. Note that the filtering capacity varies for each individual sponge specimen, highlighting the need to perform WT and MT experiments in the same aquarium to derive rigorous conclusions.

Our first observation was that it is important to use a bacterial concentration of $10^6$ cells per ml seawater as measured by flow cytometry. More ($10^7$) overwhelms the sponge leading to osculum closure and filtering cessation (**SI Figure S2**). Less ($10^5$) prevents accurate cell quantification over the course of the experiment. We used flow cytometry both to determine bacterial cell count before starting the experiments and for time course quantification (**Figure S3**).

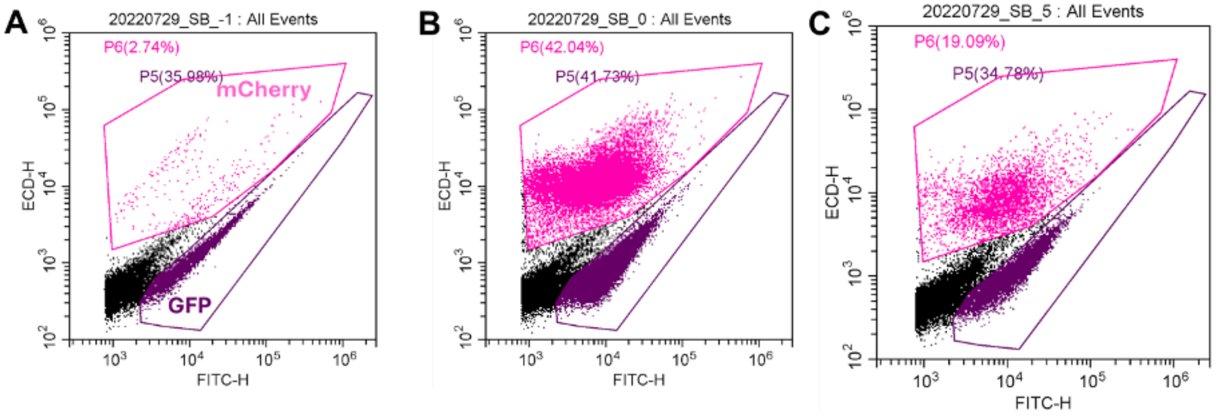


**Figure S3**. Example of flow cytometry data. Flow cytometry was used for relative cell count of *P. brasiliensis* bacteria expressing either GFP or mCherry as presented in Figure S2. (**A**) Aquarium seawater before inoculation. (**B**) Time 0 directly after inoculation. (**C**) End point. The pink and purple gates represent mCherry and GFP, respectively.

The second observation was that there was more interference in the GFP window (**Figure S3A**) due to background green fluorescence. Note the sponges are wild specimens and not axenic–no axenic sponge model has been reported. The green fluorescence interference is more apparent in the no-sponge aquaria, likely due to the growth of contaminating bacteria, which can be filtered out by the sponges in the sponge aquaria (**Figure S2B**). It was still possible to accurately compare wild-type and mutant using GFP in the sponge aquaria as we corrected for the water background (**Figure S2B**).

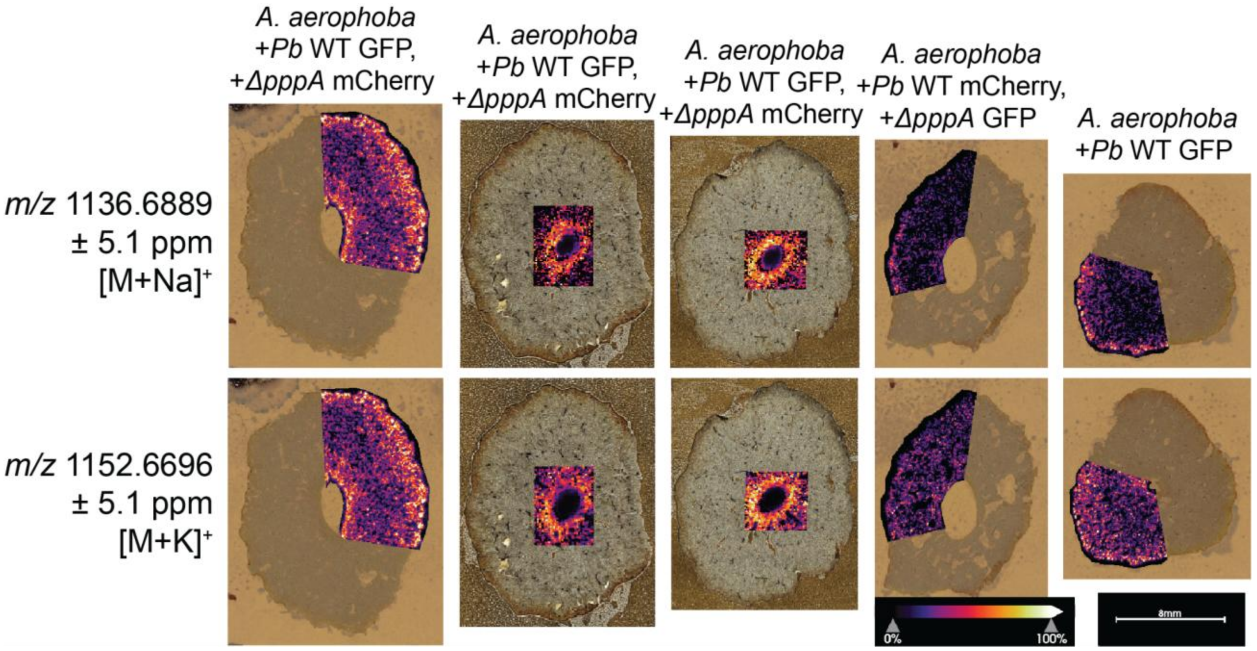


**Figure S4.** Additional MALDI-MSI fistularin-3 replicates. Both $[M+Na]^+$ and $[M+K]^+$ adducts are shown for several sponge samples. Experimental conditions shown above image.

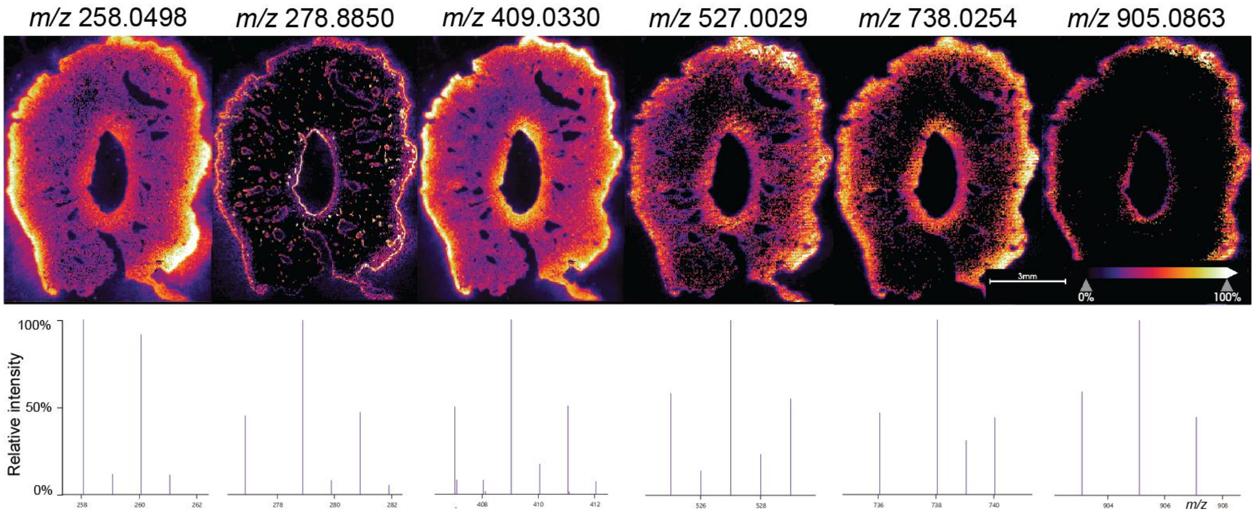


**Figure S5.** Additional brominated MALDI-MSI signals. These images are a part of the same dataset from Figure 3 of the main text. Ion images are shown along the top row, while corresponding MS isotopologue patterns are shown below each ion image. All images except the first image represent species likely containing $Br_2$, while the first image likely contains a single Br. The identity of these compounds is unknown.

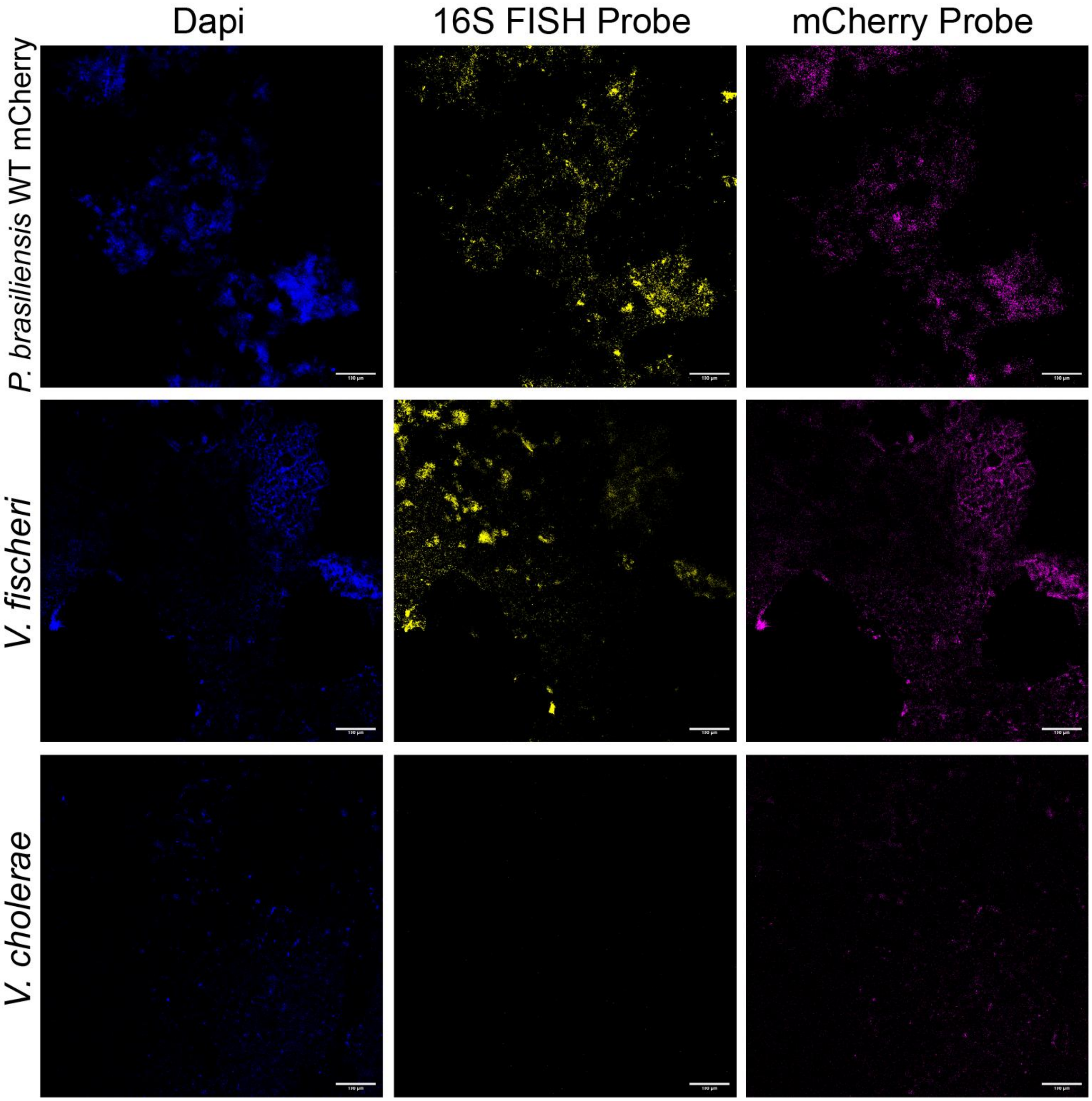


**Figure S6.** Confocal FISH images of bacterial spots. Images of *P. brasiliensis* WT mCherry, *V. fischeri*, and *V. cholerae* are shown through DAPI, the 16S FISH Probe, and the mCherry FISH probe.

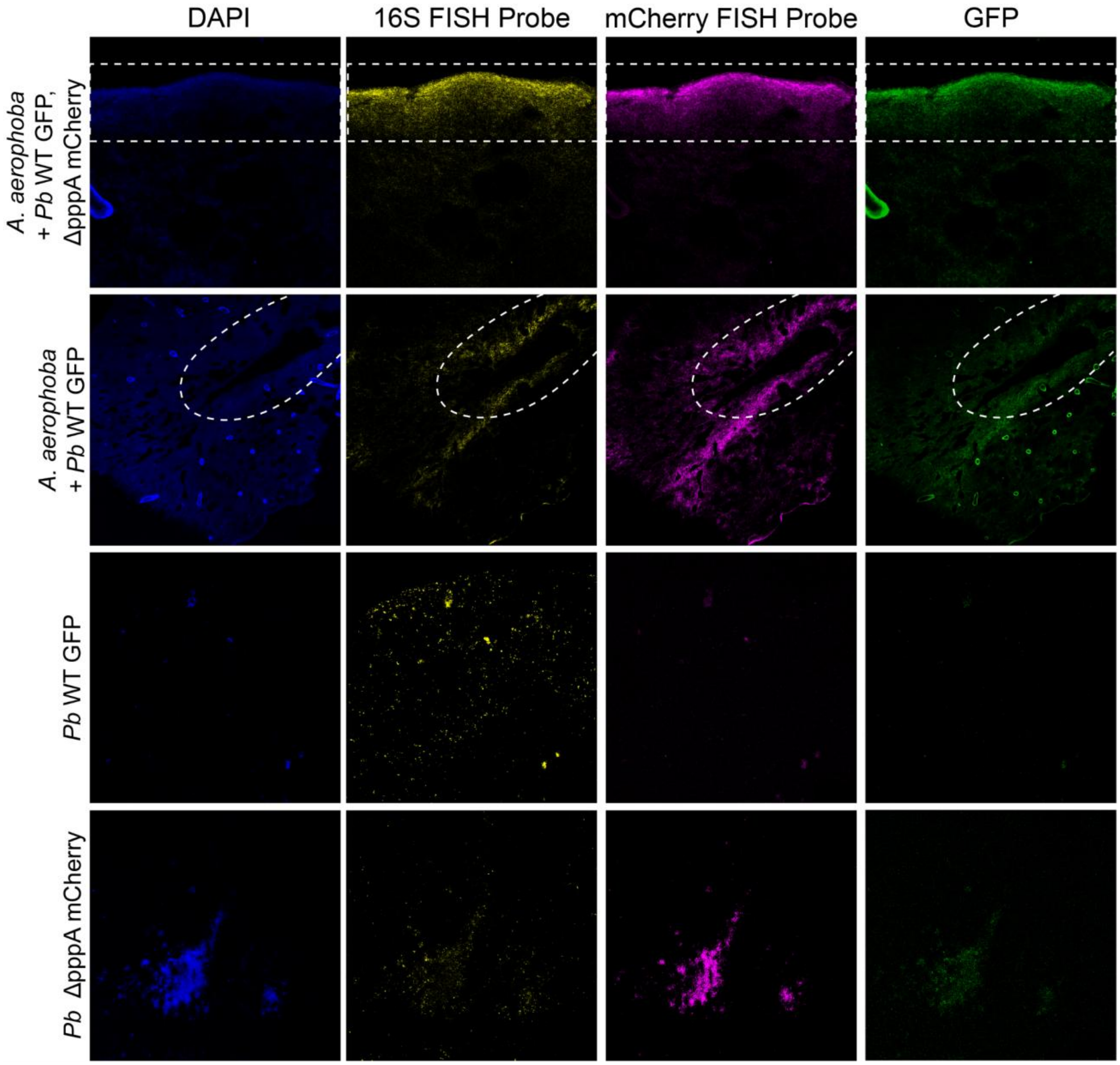


**Figure S7.** Additional confocal images of *A. aerophoba* and *P. brasiliensis.* The top row shows images of *A. aerophoba* inoculated with *P. brasiliensis* WT GFP and *ΔpppA* mCherry. The outer edge of the sponge is outlined in the white dashed box. The second row shows *A. aerophoba* inoculated with *P brasiliensis* WT GFP only. The oscule of the sponge is denoted by the white dashed line. The bottom two rows show *P. brasiliensis* WT GFP and *ΔpppA* mCherry.

| Organism | Common term | Strain | Genotype | Citation |
|---|---|---|---|---|
| *P. brasiliensis* | WT mCherry | Ab134 | pSEVA237R_Pe m7 | (2) |
| *P. brasiliensis* | WT GFP | Ab134 | pAM4891 | (2) |
| *P. brasiliensis* | Δ*pppA* mCherry | Ab134 | Δ*pppA* pSEVA237R_Pe m7 | (2) |
| *P. brasiliensis* | Δ*pppA* GFP | Ab134 | Δ*pppA* pAM4891 | (2) |
| *V. fischeri* | Biofilm down | MJM2386 | ES114 pBinK | (3) |
| *V. cholerae* | Smooth | Fy_Vc_1 | O1 El Tor A1552, wild type, Rifr | (4) |
| *P. aeruginosa* | PA01 | | | (5) |
| *B. subtilis* | 3610 | NCIB 3610 | | (6) |
| *E. coli* | K12 | | | (7) |

**Table S1.** Strain table for all organisms used in this study.

**Table S2. Gene sequences used for HCR-RNA-FISH probe design**

Pseudovibrio 16S rRNA probe, B1 amplifier, Amplifier fluorophore: 647

>16S rRNA gene sequence from Pseudovibrio Ab134 (Accession: CP074126.1:998148-999629)

AAACCTGAGAGTTTGATCCTGGCTCAGAACGAACGCTGGCGGCAGGCCTAACACAT
GCAAGTCGAACGGATCCTTCGGGATTAGTGGCAGACGGGTGAGTAACGCGTGGGAA
GCTACCTTGTGGTAGGGAACAACAGTTGGAAACGACTGCTAATACCCTATGAGCCCT
ATGGGGGAAAGATTTATCGCCATGAGATGTGCCCGCGTTAGATTAGCTAGTTGGTAA
GGTAATGGCTTACCAAGGCGACGATCTATAGCTGGTCTGAGAGGATGATCAGCCAC
ACTGGGACTGAGACACGGCCCAGACTCCTACGGGAGGCAGCAGTGGGGAATATTGG
ACAATGGGGGCAACCCTGATCCAGCCATGCCGCGTGTGTGATGACGGCCTTAGGGTT
GTAAAGCACTTTCAGCAGTGAAGATAATGACATTAACTGCAGAAGAAGCCCCGGCT
AACTTCGTGCCAGCAGCCGCGGTAATACGAAGGGGGCTAGCGTTGTTCGGAATCAC
TGGGCGTAAAGCGTACGTAGGCGGACTGATCAGTCAGGGGTGAAATCCCGGGGCTC
AACCCCGGAACTGCCTTTGATACTGTCAGTCTTGAGATCGAGAGAGGTGAGTGGAA
CTCCGAGTGTAGAGGTGAAATTCGTAGATATTCGGAAGAACACCAGTGGCGAAGGC
GGCTCACTGGCTCGATACTGACGCTGAGGTACGAAAGCGTGGGGAGCAAACAGGAT
TAGATACCCTGGTAGTCCACGCCGTAAACGATGAATGCTAGTTGTCAGGTAGCTTGC
TATTTGGTGACGCAGCTAACGCATTAAGCATTCCGCCTGGGGAGTACGGTCGCAAGA
TTAAAACTCAAAGGAATTGACGGGGGCCCGCACAAGCGGTGGAGCATGTGGTTTAA
TTCGAAGCAACGCGCAGAACCTTACCAGCCCTTGACATTTGGCGCTACATCGGGAG
ACCGATGGTTCCCTTCGGGGACGTCAGGACAGGTGCTGCATGGCTGTCGTCAGCTCG
TGTCGTGAGATGTTGGGTTAAGTCCCGCAACGAGCGCAACCCTCGCCCTTAGTTGCC
AGCATTTAGTTGGGCACTCTAGGGGGACTGCCGGTGATAAGCCGGAGGAAGGTGGG
GATGACGTCAAGTCCTCATGGCCCTTACGGGCTGGGCTACACACGTGCTACAATGGC
GGTGACAGTGGGCAGCGACCTCGCGAGAGGAAGCTAATCTCCAAAAGCCGTCTCAG
TTCGGATTGTTCTCTGCAACTCGAGAGCATGAAGTTGGAATCGCTAGTAATCGCGTA
ACAGCATGACGCGGTGAATACGTTCCCGGGCCTTGTACACACCGCCCGTCACACCAT
GGGAGTTGGTTTTACCCGAAGGCGCTGTGCTAACCGCAAGGAGGCAGGCGACCACG
GTAGGGTCAGCGACTGGGGTGAAGTCGTAACAAGGTAGCCCTAGGGGAACCTGGGG
CTGGATCACCTCCTTT

For the mCherry probe, B2 amplifier, Amplifier fluorophore: 546

>mCherry (Accession: JX560410.1:127-837)

ATGGTGAGCAAGGGCGAGGAGGATAACATGGCCATCATCAAGGAGTTCATGCGCTT
CAAGGTGCACATGGAGGGCTCCGTGAACGGCCACGAGTTCGAGATCGAGGGCGAGG
GCGAGGGCCGCCCCTACGAGGGCACCCAGACCGCCAAGCTGAAGGTGACCAAGGGT
GGCCCCCTGCCCTTCGCCTGGGACATCCTGTCCCCTCAGTTCATGTACGGCTCCAAG
GCCTACGTGAAGCACCCCGCCGACATCCCCGACTACTTGAAGCTGTCCTTCCCCGAG
GGCTTCAAGTGGGAGCGCGTGATGAACTTCGAGGACGGCGGCGTGGTGACCGTGAC
CCAGGACTCCTCCCTGCAAGACGGCGAGTTCATCTACAAGGTGAAGCTGCGCGGCA
CCAACTTCCCCTCCGACGGCCCCGTAATGCAGAAGAAGACCATGGGCTGGGAGGCC
TCCTCCGAGCGGATGTACCCCGAGGACGGCGCCCTGAAGGGCGAGATCAAGCAGAG

GCTGAAGCTGAAGGACGGCGGCCACTACGACGCTGAGGTCAAGACCACCTACAAGG
CCAAGAAGCCCGTGCAGCTGCCCGGCGCCTACAACGTCAACATCAAGTTGGACATC
ACCTCCCACAACGAGGACTACACCATCGTGGAACAGTACGAACGCGCCGAGGGCCG
CCACTCCACCGGCGGCATGGACGAGCTGTACAAGTAA
